# A shared neural code for social interaction encoding and memory in the human superior temporal sulcus

**DOI:** 10.1101/2022.10.03.510639

**Authors:** Haemy Lee Masson, Janice Chen, Leyla Isik

**Author notes:** Haemy Lee Masson and Leyla Isik, **Email:**. **Author Contributions:** H.L.M., J.C., and L.I. designed research; H.L.M. and J.C. curated data; H.L.M. analyzed data; and H.L.M., J.C., and L.I. wrote the paper.

## Abstract

Recognizing and remembering social information is a crucial cognitive skill. Neural patterns in the superior temporal sulcus (STS) support our ability to perceive others’ social interactions. However, despite the prominence of social interactions in memory, the neural basis of retrieving social interactions is still unknown. To fill this gap, we investigated the brain mechanisms underlying memory of others’ social interactions during free spoken recall of a naturalistic movie. By applying machine learning-based fMRI encoding analyses to densely labeled movie and recall data we found that STS activity patterns evoked by viewing social interactions predicted neural responses to social interaction memories. This finding suggests that the STS contains high-level conceptual, representations of social interactions, and its reactivation underlies our ability to remember others’ interactions.

## Main

Social content is a driving factor of human memory and has a profound effect on social behavior. Social cognitive brain regions, including the medial prefrontal cortex (mPFC), temporoparietal junction (TPJ), and superior temporal sulcus (STS), are fundamental to social memory. These regions are activated during social memory tasks (1) and their functional connectivity during rest predicts the quality of social memory consolidation (2, 3). Moreover, the social brain has long been implicated in general episodic memory and event retrieval. Reliable neural patterns across and within subjects in the mPFC, TPJ, and STS during event encoding are linked to better event memory (4–6). These regions are also activated during narrative free recall and the activity patterns present during event encoding are reinstated during retrieval (7). While there has been much work showing shared neural patterns across perception and memory of specific types of visual content (8), little work has been done to understand the neural basis of specific social content in memory.

Social interactions are a critical part of narrative encoding and are selectively processed in the human STS (9, 10), even during natural movie viewing when controlling for other co-varying perceptual and social features (11). Here we ask whether memory for social interactions reactivates the neural patterns evoked by perceiving social interactions in a natural movie.

Functional magnetic resonance imaging (fMRI) data were collected while participants watched a movie and then verbally described what they recalled of the movie (7). We trained a voxel-wise encoding model to link social interactions to fMRI activity during movie viewing, while accounting for other perceptual and social features in the movie. We then evaluated the model on recall data to identify brain regions with shared representations between perception and retrieval of social interactions (Fig. 1).

**Figure 1.**
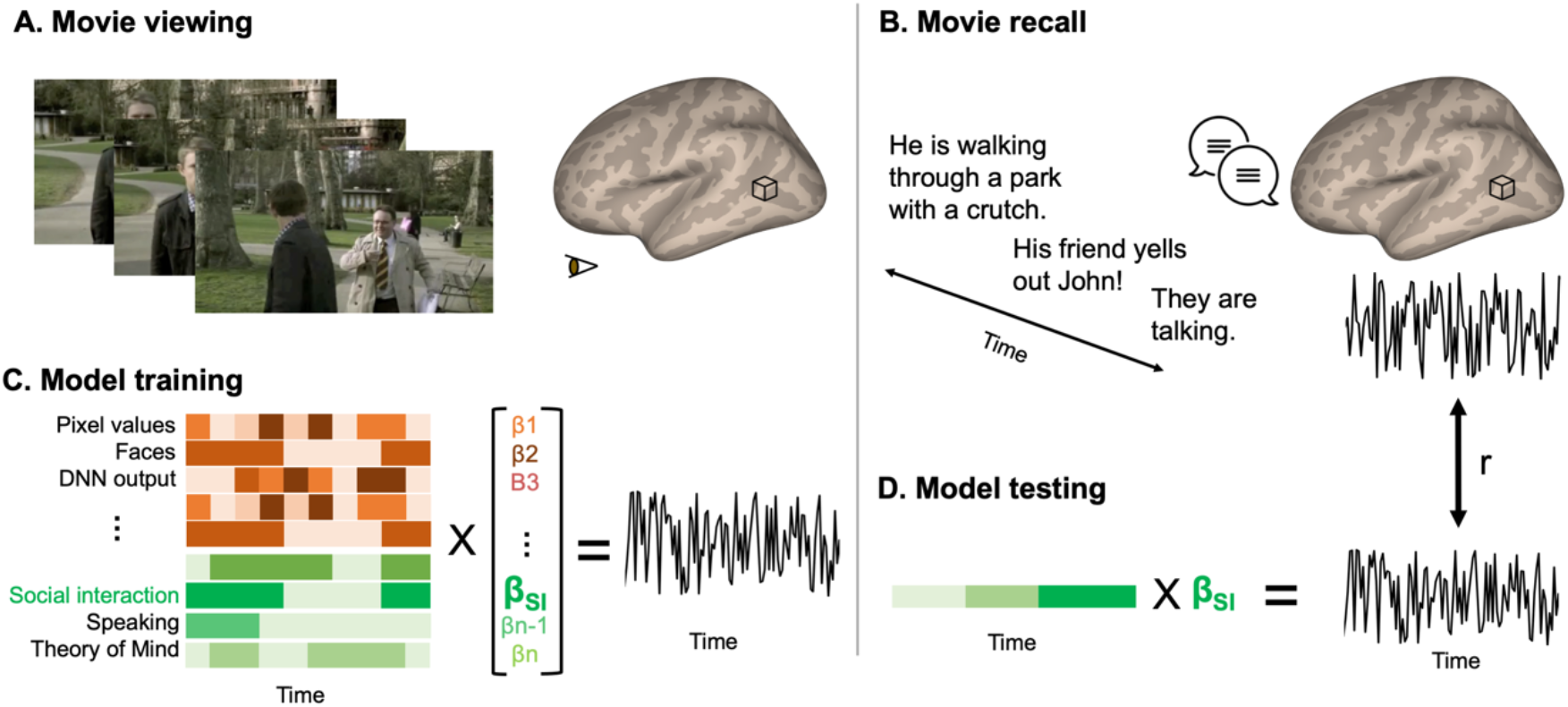
Experimental procedure and encoding framework. A) Participants watched the first episode of the Sherlock BBC series while their fMRI activity was recorded. B) In a separate scan session, participants described what they remembered about the movie during an unguided recall session. C) We trained an encoding model on the movie fMRI data. We labeled perceptual (orange) and social-affective (green) features, including the presence/absence of social interactions, at each time point in the movie. The model learned a linear mapping (beta weights) between each feature and the time-resolved data in each voxel. D) To examine the shared neural code between social interaction perception and memory, we tested the encoding model on the recall data. We labeled each time point of each participants’ spoken recall based on whether a recalled event involved social interaction (green vector). To test the encoding model, we used the beta weight specific to viewing social interactions (β_SI_) to predict brain activity evoked by social interactions during free spoken recall by multiplying it with the social interaction feature of a recalled event (green vector). The predicted brain activity was correlated with the true activity, which yielded a prediction performance score (Pearson r) assigned to each voxel.

During movie viewing, the extrastriate body area (EBA), bilateral STS, posterior medial cortex (PMC), and mPFC showed increased activation to scenes with social interactions (Fig. 2A). In contrast, scenes without social interactions activated brain areas located in visual regions across the ventral and dorsal pathways, including the inferior temporal gyrus and the inferior parietal lobe. These results confirm previous studies reporting the involvement of social brain areas, particularly the EBA (12) and STS (9–11), in social interaction perception.

**Figure 2.**
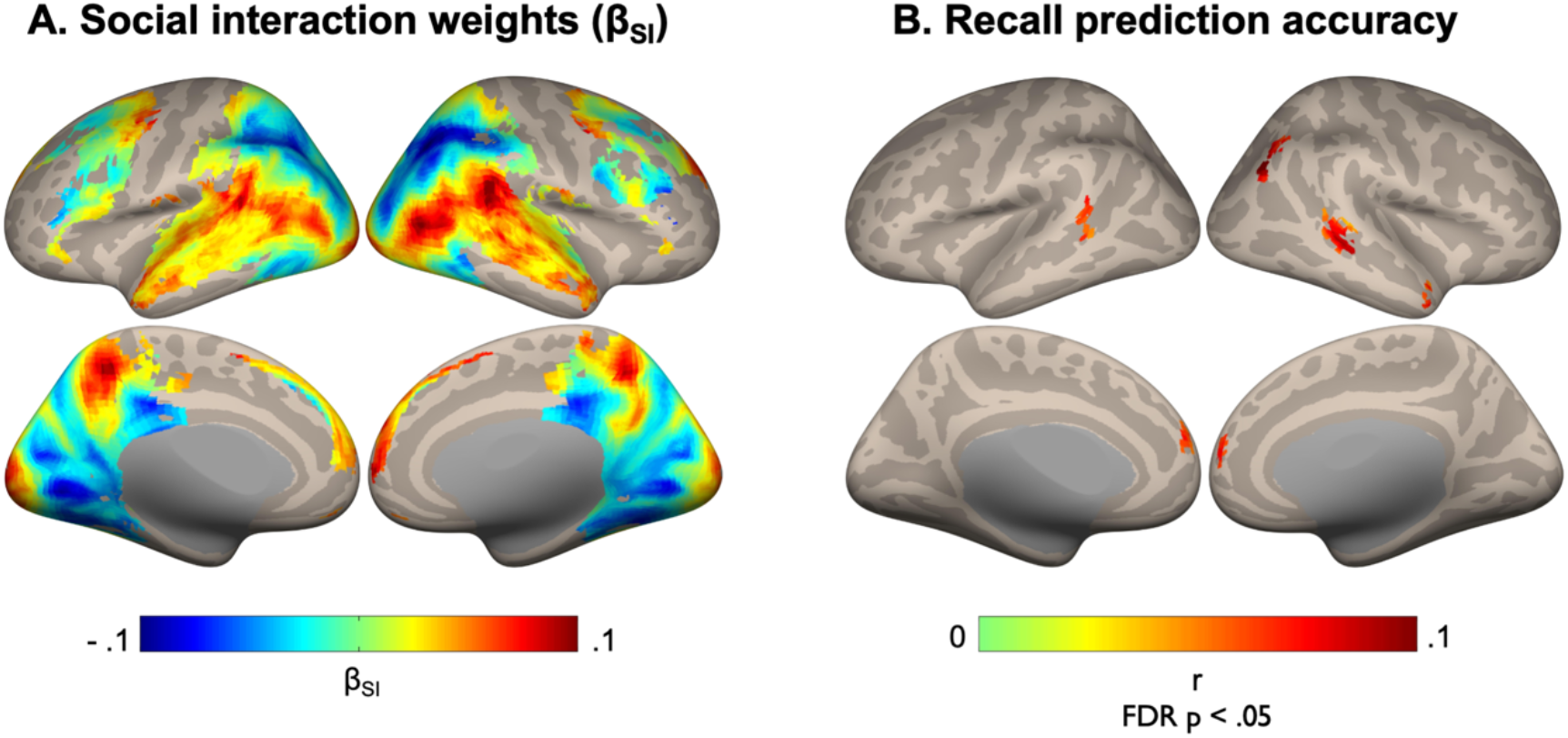
**A. Social interaction weights learned during movie viewing**. Learned beta weight for the presence of social interactions estimated from full movie viewing encoding model mapped on an inflated cortical surface. Observing social interactions (versus scenes without an interaction) elicited brain responses in EBA, STS, mPFC, and precuneus, shown yellow/red. The opposite contrast is shown in blue/purple. **B. Model prediction accuracy on recall data**. After false discovery rate (FDR) correction, the voxels that are significantly predicted (correlation, Pearson r) by social interactions in recall using the encoding model trained on movie viewing data are mapped on inflated cortices. Prediction accuracy was highest in the right STS.

During recall, participants spent on average 57% of their time describing remembered social interactions. Using the linear mapping learned to link social interactions with brain activity during movie viewing, we could accurately predict brain responses to the presence of social interactions during recall in the right temporal pole (peak accuracy found at the MNI coordinates X, Y, Z = 51, 9, −33), right inferior parietal lobe (X, Y, Z = 39, −75, 33), bilateral STS (X, Y, Z = 48, −36, −3 and −60, −48, 15), and mPFC (X, Y, Z = 6, 57, 12) (Fig. 2B). Prediction in the inferior parietal lobe was likely due to the absence of social interactions, as indicated by the negative beta weights in that region (Fig. 2A), while all other regions are likely due to the presence of social interactions. The highest prediction performance was found in the right STS for social interaction memory. Despite increased activity in the EBA during social interaction perception (Fig. 2A), we did not observe its involvement in memory retrieval.

In this study, we investigated the neural basis of memory for social interactions, and the extent to which such memory retrieval reactivates the brain regions engaged during social interaction perception. By conducting cross-validated encoding model analyses between movie viewing and free recall, we revealed that social interaction memory showed the strongest reactivation in the right STS. The posterior STS has been shown to be selectively engaged when viewing social interactions in controlled stimuli (9, 10), and this activity was replicated in our movie viewing data (Fig. 2A). Intriguingly, memory reactivation was found in a region more anterior along the mid-STS (Fig. 2B), overlapping with regions showing the strongest unique selectivity to social interactions during movie viewing (11). Our findings show that the STS is not merely involved in the perception of social interactions, but also contains amodal representations of social interactions that are engaged in the absence of any external stimulus.

Cortical reactivation for sensory memories has long been known, particularly for perceiving and remembering images and sounds (8), raising questions about the involvement of mental imagery in social interaction memory. In the current study, however, we did not find evidence of visual or auditory imagery associated with low-level perceptual features. First, a participant rarely focused on the low-level perceptual aspects of social interactions during free recall (7). Second, no reactivation was found in the visual or auditory cortex, not even in EBA, which was activated during social interaction encoding (Fig. 2A) and has been previously implicated in two body interaction perception (12). One may also question whether speech production and perception associated with self-speech during recall might have activated the STS. Given that speech production and perception do not vary systematically between recall with versus without a social interaction, it is unlikely that speech explains the STS social interaction memory prediction.

The brain areas re-activated by remembering social interactions are only a subset of those previously identified in high-level event retrieval (6, 7). By combining dense movie and recall labeling with advanced machine learning methods, we were able to link shared neural patterns between encoding and recall to specific content in narrative events. Future work is needed to understand how the neural representation of social content in the STS is integrated into general event representations.

## Materials and Methods

We analyzed fMRI data and audio transcripts from the original study (7). The Princeton University Institutional Review Board approved the original study. All participants provided their written informed consent before the experiment. A selective overview of methods is included. More detailed descriptions can be found in two studies: participants, experimental procedure, fMRI data acquisition and preprocessing (7); encoding analysis methods and movie annotations (11).

### fMRI data sources

fMRI data consisted of 3 functional runs from 16 participants, with 2 runs from movie viewing and 1 from movie recall (Figure 1). 16 participants viewed the first episode of the Sherlock BBC series and then freely recalled scenes in the scanner. The duration of movie viewing was about 45 mins in total, and the duration of the recall varied across participants.

### fMRI data acquisition and preprocessing

In each functional run, whole-brain images (27 slices, voxel size = 4 x 3 x 3 mm^3^) were collected on a 3T Siemens Skyra scanner with a 20-channel head coil. An echo-planar T2* weighted sequence was used with the acquisition parameters of repetition time (TR) = 1500 ms, echo time = 28 ms, flip angle = 64°, and field of view = 192 x 192 mm^2^.

Preprocessing included correction of slice-timing and head movements, detrending of blood-oxygen-level-dependent (BOLD) signal, temporal high-pass filtering (140 s cut off), normalization to a Montreal Neurological Institute (MNI) space with voxels re-sampled to 3 x 3 x 3 mm^3^, and smoothing with a 6-mm full width at half maximum Gaussian kernel. Preprocessed time series data were z-score standardized and shifted by 3 TRs from the onset to correct for the hemodynamic delay.

### Audio transcripts and feature annotation

Audio recordings made during free recall were transcribed in the original study (7). We labeled the presence versus absence of social interactions at each time point of each participant’s recall. To do that, first, we divided transcripts into sentences. Next, using a 1-5 scale, two human annotators judged whether each sentence described a social interaction (*Is a participant talking about a social interaction? 1-strongly disagree, 5 strongly agree*). Ratings were averaged and binarized (scores below 3 were labeled as non social interaction; scores above 3 were labeled as a social interaction). Note that judgments from the two annotators were strongly correlated (r = 0.8). This binarized feature was used as a predictor in our regression model (Fig. 1D, green vector).

### Voxel-wise encoding modeling

Using cross-validated encoding analyses (Fig. 1), we learned a linear mapping between the presence of social interactions (in addition other labeled perceptual and social movie features) and brain activity during movie viewing (model training), and then examined whether the same linear mapping could link the presence of social interactions to brain activity during recall (model testing). Specifically, we trained an encoding model consisting of 16 perceptual and social-affective features of the movie, including the presence of social interaction between people in the movie (Fig. 1C). A beta weight for each feature and voxel was estimated during training to link movie features to fMRI responses during movie viewing (11). We used linear banded ridge regression to account for high-dimensional features (including the output of a deep neural network) and multiple collinearities between features. To remove unreliable voxels, we excluded voxels outside of the brain mask created based on inter-subject correlation values in our previous study (11). In brief, this mask only included voxels showing shared stimuli-evoked responses across participants during movie viewing. During testing, the beta weights learned for viewed social interactions were multiplied by the social interaction recall feature (the presence vs absence of social interactions during recall) to predict the withheld BOLD responses recorded during movie recall (Fig. 1D). We calculated the prediction performance scores based on the similarity between the true BOLD and estimated BOLD signals across participants. Statistical inference was made with a sign permutation test (5000 iterations). One-tailed P values were calculated and adjusted with false discovery rate (FDR) correction. Group averaged prediction performance was thresholded at P _FDR_ < 0.05 and visualized on the cortical surface using the CONN software (13). More details on the voxel-wise encoding analyses are available in our previous study (11).

## Data availability

Preprocessed fMRI data are available at (https://dataspace.princeton.edu/handle/88435/dsp01nz8062179). Codes are available at (https://github.com/haemyleemasson/voxelwise_encoding).

## Acknowledgments

This work was supported with funds from The Clare Boothe Luce Program for Women in STEM. We thank Lucy Chang and Qing Lu or assisting with feature annotation.

## References

1. M. L. Meyer, E. Collier, Theory of minds: managing mental state inferences in working memory is associated with the dorsomedial subsystem of the default network and social integration. Social cognitive and affective neuroscience 15, 63–73 (2020).

2. E. Collier, M. L. Meyer, Memory of Others’ Disclosures Is Consolidated during Rest and Associated with Providing Support: Neural and Linguistic Evidence. Journal of cognitive neuroscience 32, 1672–1687 (2020).

3. M. L. Meyer, L. Davachi, K. N. Ochsner, M. D. Lieberman, Evidence That Default Network Connectivity During Rest Consolidates Social Information. Cerebral cortex (New York, N.Y.: 1991) 29, 1910 (1920).

4. U. Hasson, O. Furman, D. Clark, Y. Dudai, L. Davachi, Enhanced Intersubject Correlations during Movie Viewing Correlate with Successful Episodic Encoding. Neuron 57, 452–462 (2008).

5. E. Simony, et al., Dynamic reconfiguration of the default mode network during narrative comprehension. Nature Communications 2016 7:1 7, 1–13 (2016).

6. C. Baldassano, et al., Discovering Event Structure in Continuous Narrative Perception and Memory. Neuron 95, 709–721.e5 (2017).

7. J. Chen, et al., Shared memories reveal shared structure in neural activity across individuals. Nature Neuroscience 20, 115–125 (2017).

8. M. Wheeler, S. Petersen, R. Buckner, Memory’s echo: Vivid remembering reactivates sensory-specific cortex. Proceedings of the National Academy of Sciences 97, 11125–11129 (2000).

9. L. Isik, K. Koldewyn, D. Beeler, N. Kanwisher, Perceiving social interactions in the posterior superior temporal sulcus. Proceedings of the National Academy of Sciences of the United States of America 114 (2017).

10. J. Walbrin, P. Downing, K. Koldewyn, Neural responses to visually observed social interactions. Neuropsychologia 112, 31–39 (2018).

11. H. Lee Masson, L. Isik, Functional selectivity for social interaction perception in the human superior temporal sulcus during natural viewing. NeuroImage 245, 118741 (2021).

12. E. Abassi, L. Papeo, The representation of two-body shapes in the human visual cortex. Journal of Neuroscience 40, 852–863 (2020).

13. S. Whitfield-Gabrieli, A. Nieto-Castanon, Conn: a functional connectivity toolbox for correlated and anticorrelated brain networks. Brain connectivity 2, 125–141 (2012).

